# Retrograde transport of mannose-6-phosphate receptor depends on several sorting machineries as analyzed by sulfatable nanobodies

**DOI:** 10.1101/2019.12.21.885939

**Authors:** Dominik P. Buser, Martin Spiess

## Abstract

Retrograde protein transport from the cell surface and endosomes to the trans-Golgi network (TGN) is essential for membrane homeostasis in general and for the recycling of mannose-6-phosphate receptors (MPRs) for sorting of lysosomal hydrolases in particular. Several different sorting machineries have been implicated in retrieval from early or late endosomes to the TGN, mostly for the cation-independent MPR (CIMPR), mainly by analysis of steady-state localization and by interaction studies. We employed a nanobody-based sulfation tool to more directly determine transport kinetics from the plasma membrane to the TGN – the site of sulfation – for the cation-dependent MPR (CDMPR) with and without silencing of candidate machinery proteins. The clathrin adaptor AP-1 that operates bidirectionally at the TGN-to-endosome interface, which had been shown to cause reduced sulfation when rapidly depleted, produced hypersulfation of nanobodies internalized by CDMPR upon long-term silencing, reflecting accumulation in the TGN. In contrast, knockdown of retromer (Vps26), epsinR, or Rab9 reduced CDMPR arrival to the TGN. No effect was observed upon silencing of TIP47. Most surprisingly, depletion of the GGA (Golgi-localized, γ-adaptin ear-containing, Arf-binding) proteins inhibited retrograde transport rather than TGN exit. This study illustrates the usefulness of derivatized, sulfation-competent nanobodies to analyze retrograde protein transport to identify the contributions of different machineries.

## Introduction

Retrograde transport of proteins from the cell surface and endosomes to the trans-Golgi network (TGN) is critical for membrane homeostasis and to retrieve components of anterograde transport machineries. Proteins recycled back to the TGN comprise transport receptors for lysosomal hydrolases, processing enzymes, SNAREs (soluble N-ethylmaleimide-sensitive fusion factor attachment receptors), nutrient transporters, and a subset of other intracellular transmembrane proteins with diverse functions (Bonifacino and Rojas, 2006; Johannes and Popoff, 2008). In addition, extracellular bacterial and plant toxins exploit the retrograde transport machineries of target cells. One of the most thoroughly studied cargoes retrieved from endosomes to the TGN are the cation-dependent and -independent mannose-6-phosphate receptors (CDMPR and CIMPR), involved in efficient anterograde transport of lysosomal acid-hydrolases from the TGN to endosomes (Ghosh et al., 2003a). Following cargo unloading in the mildly acidic endosomal environment, MPRs are recycled to the TGN apparently by several retrograde transport machineries from both early/recycling and late endosomes (Bonifacino and Rojas, 2006; McKenzie et al., 2012; Pfeffer, 2009; Sandvig and van Deurs, 2002).

The most prominent sorting machinery reported to mediate transport of MPRs to the TGN is the retromer complex, a pentameric protein assmbly comprising Vps (vacuolar protein sorting) and SNX-BAR (sorting nexin-Bin/Amphiphyisin/Rvs) subunits (Bonifacino and Hurley, 2008; Gallon and Cullen, 2015; Seaman, 2012). The core complex, termed retromer, consists of the heterotrimer Vps26-Vps29-Vps35 that transiently associates with the tubulation subcomplexes composed of SNX1 or SNX2, and SNX5 or SNX6. Other SNX-BAR proteins, including SNX3, have been shown to mediate endosome-to-TGN transport of WLS (Belenkaya et al., 2008; Harterink et al., 2011). The precise sites from which retromer complexes operate remain to be defined: Vps35 of retromer was shown to be recruited by Rab7a, a marker of late endosomes, but SNX-BAR components of the multimeric complex bind via their Phox homology (PX) domain to phosphatidylinositol 3-phosphate (PI3P), a phospholipid enriched on early endosomes (Carlton et al., 2004; Carlton et al., 2005; Cozier et al., 2002; Rojas et al., 2008; Seaman et al., 2009). It was thus proposed that MPR sorting by retromer complex is a progressive process coupled to endosomal maturation during the Rab5-to-Rab7 switch (Rojas et al., 2008). Some uncertainty also exists about cargo recognition by Vps and/or SNX-BAR subunits. Previously, it was reported that Vps subunits serve as cargo adaptors for the cytoplasmic domain of CIMPR (Cui et al., 2019; Fjorback et al., 2012; Lucas et al., 2016; Nothwehr et al., 2000; Seaman, 2007; Suzuki et al., 2019). Two recent independent studies rather suggest that SNX1/2 and SNX5/6 mediate cargo recognition and retrieval of CIMPR (Kvainickas et al., 2017; Simonetti et al., 2017). Their results not only showed that SNX-BAR dimers associate with a WLM motif in the cytoplasmic tail of CIMPR, but also that Vps35 depletion, unlike depletion of SNX-BARs, did not cause receptor misdistribution from juxtanuclear to peripheral compartments. This observation is in disagreement with previous results showing a prominent mislocalization of CIMPR to endosomes upon silencing of Vps26 or Vps35 (Arighi et al., 2004; Seaman, 2004).

A further pathway for retrograde transport of MPRs was described to involve Rab9 and the adaptor TIP47 (tail-interacting protein of 47 kDa). Using a cell-free system, it was shown that Rab9 recruits TIP47 to late endosomes and that interference with GTPase-effector function resulted in severe impairment of transport of MPRs (Diaz and Pfeffer, 1998; Lombardi et al., 1993). However, TIP47 was since identified to be a component of lipid droplets involved in their biogenesis (Bulankina et al., 2009) and an additional role in retrograde transport was not independently reproduced (Medigeshi and Schu, 2003).

Another mechanism for MPR retrieval to the TGN involves clathrin-coated vesicles (CCVs) with the adaptor protein (AP)-1 complex and/or epsinR. AP-1 has a well-established role in anterograde transport of MPRs from the TGN to endosomes in cooperation with GGA (Golgi-localized, γ-adaptin ear-containing, Arf-binding) proteins (Doray et al., 2002; Ghosh and Kornfeld, 2004; Sanger et al., 2019). Unlike retromer complexes, AP-1 has a dual distribution both at the TGN and on early endosomes (Le Borgne et al., 1996; Meyer et al., 2000; Seaman et al., 1996). Inactivation of AP-1 was found to result in a dispersed MPR localization pattern towards the periphery of cells (Meyer et al., 2000; Robinson et al., 2010), similar to the phenotype of retromer complex inactivation, suggesting a role in retrograde transport. EpsinR, an interactor of AP-1, was also shown to have a role in CIMPR retrieval to the TGN (Hirst et al., 2004; Saint-Pol et al., 2004), although receptor distribution did not significantly change in epsinR-depleted cells. It is puzzling, why epsinR and AP-1 have different effects on MPR localization when depleted, while they seem to depend on each other for incorporation into CCVs, depletion of one reducing the CCV content of the other (Hirst et al., 2015; Hirst et al., 2004).

GGAs also localize to both TGN and endosomes (Boman et al., 2000; Ghosh et al., 2003b). Yet, they have been implicated mainly in anterograde transport. Rapid depletion of GGA2 by knocksideways specifically depleted lysosomal hydrolases and their receptors (MPRs and sortillin) from CCV contents, while knocksideways of AP-1 affected also a number of SNAREs and additional membrane proteins (Hirst et al., 2012). Depletion of lysosomal hydrolases from CCVs was more efficient upon inactivation of GGA2 and depletion of their receptors more efficient upon inactivation of AP-1. This result suggested a role of GGAs primarily in anterograde transport and of AP-1 in both directions.

To analyze retrograde transport machinery, most studies used fluorescence microscopy to monitor changes in MPR localization relative to TGN-resident or endosomal markers by statistical steady-state image analysis. A few laboratories imaged antibody uptake to follow retrograde transport from the cell surface to the TGN (e.g., Breusegem and Seaman, 2014), while Johannes and colleagues employed sulfation as a specific modification of the trans-Golgi/TGN to probe Golgi arrival of Shiga toxin B-chain (STxB) tagged with a sulfation motif or of antibodies derivatized with sulfatable peptides (Popoff et al., 2009; Saint-Pol et al., 2004). However, the disadvantage of conventional divalent antibodies is that they are rather large and can crosslink their antigens and thus potentially alter their trafficking. In contrast, monomeric protein binders, such as nanobodies, are monovalent and small. Nanobodies are easily derivatized with sequence tags and fluorescent or enzymatic protein domains and can be produced in bacteria.

To study retrograde traffic, we have previously established a versatile toolkit of functionalized anti-GFP nanobodies (Buser et al., 2018; Buser and Spiess, 2019). In particular, we generated nanobodies containing tyrosine sulfation (TS) sites to monitor their arrival in the trans-Golgi/TGN. Using cell lines stably expressing EGFP-CDMPR or -CIMPR, we determined the transport kinetics of these receptors from the cell surface to the TGN. In addition, we used the knocksideways system developed by Robinson et al. (2010) to analyze the contribution of AP-1 upon rapid depletion. The system is based on rapamycin-induced heterodimerization between the γ-subunit of AP-1 fused to FKBP12 (FK506-binding protein of 12 kDa) and the FKBP–rapamycin-binding domain (FRB) of mTOR (mammalian target of rapamycin) anchored in the outer mitochondrial membrane as a trap (Mitotrap). Upon rapid inactivation of AP-1, a robust reduction of sulfation kinetics by appoximately on third was observed, confirming a significant contribution of AP-1/clathrin-coated vesicles in retrograde transport of MPRs (Buser et al., 2018).

In the present study, we applied this tool of sulfatable nanobodies to analyze plasma membrane-to-TGN transport kinetics of CDMPR in order to define the contribution of individual different sorting machineries in parallel on retrograde transport in living cells. We could confirm retrograde transport activity of retromer and epsinR as well as an involvment of Rab9, but not of TIP47. Unexpectedly, silencing of GGA1–3 also reduced CDMPR arrival at the TGN, suggesting a role of GGAs in endosome-to-TGN rather than anterograde transport.

## Results

### Functionalized nanobodies to analyze retrograde transport of CDMPR in cells depleted of candidate machineries

To study retrograde traffic to intracellular compartments including the TGN, we have previously established a versatile toolkit of functionalized anti-GFP nanobodies (Buser et al., 2018; Buser and Spiess, 2019). They can be used to label GFP-tagged proteins of interest at the cell surface and follow their route to endosomes, the TGN, and back to the plasma membrane. Here, we employed anti-GFP nanobodies (VHH, variable heavy-chain domain of heavy-chain–only antibody) modified with a hexahistidine tag for purification, a T7 and an HA tag for immunodetection, a biotin acceptor peptide for biotinylation, and sequences conferring tyrosine sulfation (VHH-2xTS) or red fluorescence (VHH-mCherry) (Fig. 1A). These functionalized nanobodies were bacterially expressed and isolated to high purity, and shown to be efficiently immunodetected using epitope tag antibodies or streptavidin-HRP (Fig. 1B).

**Figure 1.**
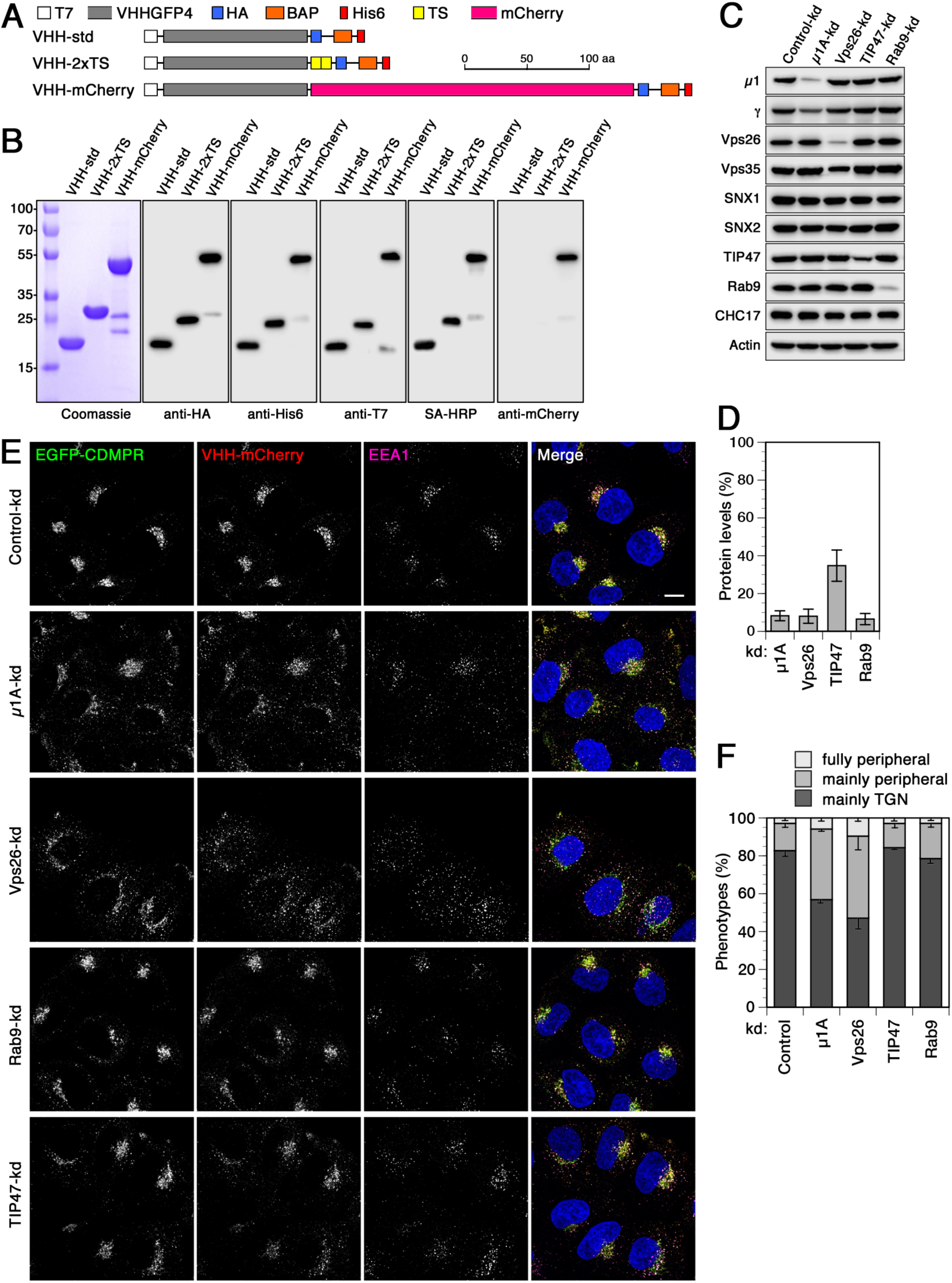
Functionalized nanobodies to analyze retrograde transport of CDMPR upon silencing of AP-1, retromer, Rab9, or TIP47. (**A**) Schematic representation of the functionalized nanobodies. The standard nanobody (VHH-std) consists of the GFP-specific VHH domain, T7 and HA epitope tags, a biotin acceptor peptide (BAP), and a hexahistidine (His6) purification tag. Other nanobodies in addition contain two tyrosine sulfation sequences (VHH-2xTS) or mCherry (VHH-mCherry). Scale bar in amino acids (aa). (**B**) Bacterially expressed and purified nanobodies (30 *µ*g) were analyzed by SDS-gel electrophoresis and Coomassie staining (left). Immunoblot analysis of nanobodies (10 ng) with antibodies against the HA, His6, T7, or mCherry epitopes, or with streptavidin-HRP (SA-HRP). Marker proteins with molecular weights in kDa are shown on the left. As previously reported (Buser et al., 2018; Buser and Spiess, 2019), mCherry-containing nanobodies are slightly susceptible to clipping between the VHH and mCherry domains. (**C**) HeLa cells were transfected with non-targeting siRNA or siRNAs targeting μ1A-adaptin, Vps26, TIP47, or Rab9. Three days after transfection, cells were subjected to immunoblot analysis with antibodies against the indicated proteins. (**D**) To determine the knockdown (kd) efficiency, the residual protein was quantified in percent of the value after control-kd (mean and standard deviation of three independent experiments). (**E**) HeLa cells stably expressing EGFP-CDMPR were depleted of μ1A-adaptin, Vps26, TIP47, or Rab9 as in (C). Cells were incubated for 1 h at 37°C with full medium containing 5 *µ*g/ml VHH-mCherry (∼0.1 *µ*M), fixed, stained for EEA1 and nuclei (DAPI, blue), and imaged by fluorescence microscopy. Bar: 10 *µ*m. (**F**) Quantitation of the percentage of cells displaying the CDMPR localization phenotypes “mainly TGN”, “mainly peripheral”, or “fully peripheral” as in Simonetti et al., 2017 and Wassmer et al., 2007. For each condition, random frames with a total of 136–140 cells were scored from three independent experiments.

A well-established phenotype of retrograde transport deficiency on MPR traffic is the redistribution of the receptor from juxtanuclear to more peripheral compartments. To analyze the contribution of individual MPR retrieval routes to the TGN, we depleted machinery components of the AP-1/clathrin-, the retromer-, and the Rab9/TIP47-dependent pathways by RNA interference using well established siRNAs. Inactivation of the AP-1 complex was achieved by depleting the μ1A-subunit of the heterotetrameric adaptor complex (Hirst et al., 2005; Hirst et al., 2003; Hirst et al., 2009). The retromer complex was inactivated by silencing the subunit Vps26 of the cargo-selective complex (CSC) (Popoff et al., 2009; Popoff et al., 2007). Rab9 and TIP47, which do not form a stable complex, were knocked down separately (Bulankina et al., 2009; Ganley et al., 2004; Kucera et al., 2016; Reddy et al., 2006). All these proteins could be robustly depleted by >85% (Fig. 1C and D), except TIP47 which was consistently reduced by ∼65%, similar to the depletion efficiencies for TIP47 in the literature where they had been reported to produce clear effects on MPR traffic in vitro and in vivo (Diaz and Pfeffer, 1998; Ganley et al., 2004). As previously observed, depletion of μ1A or Vps26 caused a concomitant reduction of its complex partners γ-adaptin or Vps35, respectively (Fig. 1C) (Arighi et al., 2004; Meyer et al., 2000), while the retromer-associated SNX-BAR proteins SNX1 and SNX2 were not affected upon depletion of Vps26 (Arighi et al., 2004; Rojas et al., 2007). Depletion of components of one pathway did not affect expression levels of proteins associated with other retrograde transport routes to the TGN (Fig. 1C).

We silenced the above machinery components in HeLa cells stably expressing EGFP-CDMPR and analyzed its steady-state localization by fluorescence microscopy (Fig. 1E) to test for mislocalization to endosomal compartments, a well-documented phenotype thought to result from defective endosome-to-TGN retrieval (Arighi et al., 2004; Hirst et al., 2012; Meyer et al., 2000; Popoff et al., 2009; Robinson et al., 2010; Seaman, 2004). In addition, we added VHH-mCherry nanobodies to the cells for 1 h before fixation to specifically detect the mature pool of the receptor cycling between surface, endosomes, and TGN (similar to previous antibody uptake experiments (Meyer et al., 2000; Robinson et al., 2010)). To measure the extent of mislocalization of EGFP-CDMPR to peripheral compartments, we used a semi-quantitative approach classifying the CDMPR staining patterns of individual cells as “mainly TGN”, “mainly peripheral”, and “fully peripheral” (Fig. 1F) as previously applied by Cullen and colleagues (Simonetti et al., 2017; Wassmer et al., 2007). Knockdown of AP-1 or retromer caused a pronounced shift of both receptor and imported nanobody from the TGN to peripheral punctae as compared to control cells. We observed an approximately 3-fold increase in peripherally dispersed MPR-nanobody distributions in AP-1- and Vps26-depleted cells (Fig. 1E and F), in agreement with previous analyses (e.g., (Wassmer et al., 2007). In contrast, MPR localization was not significantly affected upon depletion of TIP47 or Rab9.

At the same time, knocking down any of these four components did not significantly alter the plasma membrane levels of EGFP-CDMPR, as was assessed by nanobody binding only to the surface receptors at 4°C (Fig. S1). As a positive control, depletion of clathrin heavy chain (CHC17), which is required for clathrin-mediated endocytosis, produced the expected clear increase in surface EGFP-CDMPR. To make sure that depletion of potential machinery components do not generally affect sulfation efficiency, on which our assay critically depends, HeLa cells stably expressing a sulfatable form of the secretory protein ✓1-protease inhibitor (A1Pi) were labeled with [^35^S]sulfate. No change in sulfation of A1Pi was observed for any of these protein knockdowns (Fig. S2).

### Retrograde transport of CDMPR to the TGN is affected by depletion of Rab9 or Vps26, but not of TIP47

To more directly, more sensitively, and more quantitatively assay endosome-to-TGN transport in control and knockdown cells, we employed VHH-2xTS nanobodies containing sites for tyrosine sulfation. This allows us to correlate appearance of nanobody sulfation with TGN arrival and residence time in this compartment. Cells stably expressing EGFP-CDMPR were silenced for one of the candidate retrograde machinery components or, as a control, transfected with non-targeting siRNA. The cells were then incubated with media containing VHH-2xTS for up to 75 min while labeling with [^35^S]sulfate. In control cells, nanobody binding to EGFP-CDMPR and uptake reached its maximum within little more than 30 min and 50% after about 15 min (Fig. 2A and B, open squares). Sulfation started only after a lag time of ∼15 min and had not yet reached saturation after 75 min (Fig. 2B, filled squares), in full agreement with our previous report (Buser et al., 2018). The difference between uptake and sulfation curves reflects the time of transport to the TGN.

**Figure 2.**
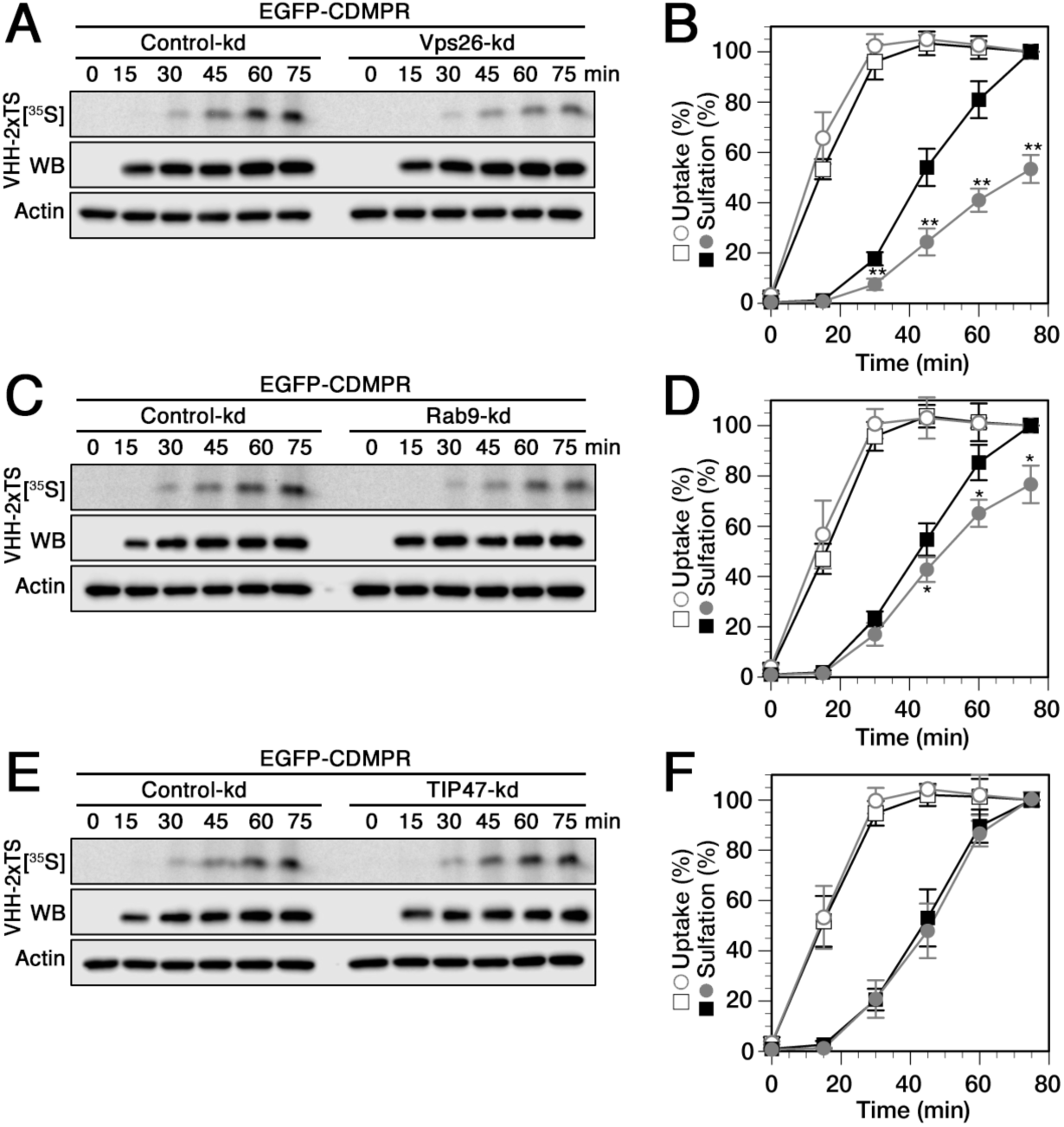
Changes in retrograde transport kinetics of CDMPR to the TGN upon silencing Vps26, Rab9, or TIP47. Cells stably expressing EGFP-CDMPR were transfected with non-targeting siRNA (control-kd) or with siRNA silencing expression of Vps26 (**A**), Rab9 (**C**), or TIP47 (**E**) as described in Fig.1. The cells were labeled with [^35^S]sulfate for up to 75 min in the presence of 2 μg/ml VHH-2xTS. The nanobodies were isolated by Ni/NTA beads and subjected to SDS-gel electrophoresis followed by Western-blot (WB) analysis (anti-His6) and autoradiography ([^35^S]). In parallel, aliquots of the cell lysates were immunoblotted for actin as a control for the amount of cells used. Three independent experiments as shown in panels A, C, and E were quantified in panels **B, D**, and **F**, respectively, and presented as the percentage of the value of control-kd cells after 75 min (mean and standard deviation of three independent experiments; two-sided Student’s *t*-test: *, p < 0.05; **, p < 0.01). Control-kd are shown as black squares and target-kd as gray circles; uptake as open symbols, sulfation as filled symbols.

The most thoroughly analyzed sorting machinery in endosome-to-TGN retrieval of MPRs is the canonical retromer complex (Arighi et al., 2004; Burd and Cullen, 2014; Seaman, 2004; Wassmer et al., 2007). Knocking down Vps26 as a core component of retromer caused a significant reduction of the rate of sulfation, indicating only ∼50% of nanobody transport to the TGN after 75 min, while uptake was not considerably affected (Fig. 2A and B). These results support a contribution of Vps26 in retrograde transport of CDMPR, confirmings previous observations by different laboratories for CIMPR using immunofluorescence and antibody uptake assays (Arighi et al., 2004; McKenzie et al., 2012; Popoff et al., 2009; Popoff et al., 2007; Seaman, 2004; Seaman, 2007). Notably, we obtained a similar extent of transport impairment for EGFP-CDMPR as previously reported for STxB using single-time point sulfation experiments (Popoff et al., 2009; Popoff et al., 2007).

Upon depletion of Rab9, nanobody uptake remained unaffected both in kinetics and in extent, whereas sulfation was modestly, but significantly reduced (Fig. 2C and D). This suggests that Rab9 contributes directly or indirectly to retrograde transport from endosomes to the compartment of sulfation. This effect is consistent with previous observations based on image analysis and antibody uptake for chimeric CIMPR and furin (Chia et al., 2011; Seaman et al., 2009). In these previous reports, transport to the TGN was reduced by up to 50% upon Rab9 knockdown, while we could only observe a reduction in signal of ∼25% after 75 min, a difference that might be due to the method used. Interestingly, while retrograde transport of EGFP-CDMPR to the TGN was affected in kinetics by Rab9 depletion, the steady-state localization of the receptor was barely affected (Fig. 1E and F), reflecting that kinetic assays are required to directly study endosome-to-TGN transport. In contrast to Ganley et al. (Ganley et al., 2004; Ganley et al., 2008), who observed increased lysosomal turnover of MPR in Rab9-depeleted cells, we could not detect any reduction of VHH-2xTS internalized piggyback by EGFP-CDMPR (Fig. 2C and D) or of the receptor itself in the knockdown cells (Fig. S1).

Since TIP47 was proposed to mediate Rab9-dependent MPR transport, knockdown should produce a similar effect as Rab9 depletion. However, kinetics of nanobody uptake and sulfation by EGFP-CDMPR remained unchanged in TIP47 knockdown cells (Fig. 2E and F). Our sulfation experiments add an additional method to those used previously to evaluate retrograde transport kinetics of CDMPR with and without TIP47, again with a negative result.

### AP-1 contributes to both retrograde and anterograde transport of CDMPR

To investigate the contribution of AP-1 in this process, we have previously analyzed the role of AP-1 in CD- and CIMPR transport to the TGN by rapid inactivation of AP-1 using knocksideways (Buser et al., 2018). Rapid depletion showed a reduction of approximately one third in the rate of sulfation, demonstrating a significant contribution of AP-1/clathrin in endosome-to-TGN transport of the MPRs. To test the outcome with AP-1 silencing in the same manner as applied to analyze the contribution of the other potential machineries above, we also performed the transport assay upon siRNA knockdown of *µ*1A.

Surprisingly, we did not observe a reduction of the kinetics and the extent of nanobody sulfation as expected from the more peripheral steady-state distribution of CDMPR in long-term AP-1-depleted cells. Instead, we found, after a similar lag phase as in all previous conditions, an approximately 2.5-fold increase in rate and extent of sulfation (Fig. 3A and B). If this reflected transport directly, it would indicate increased retrograde transport activity by other mechanisms that even strongly overcompensated the loss of AP-1/clathrin-mediated transport. Yet, no increase in the levels of other machineries were detectable in *µ*1A knockdown cells (Fig. 1C and Fig. S1) and no increase in general tyrosine sulfation (Fig. S2). The strong sulfation signal is also not the result of increased uptake of VHH-2xTS, since the signals of cell-associated nanobody after loading at 37°C to steady-state or after binding only to cell-surface EGFP-CDMPR at 4°C were not increased in AP-1 knockdown compared to control cells (Fig. 3A and B, and Fig. S1).

**Figure 3.**
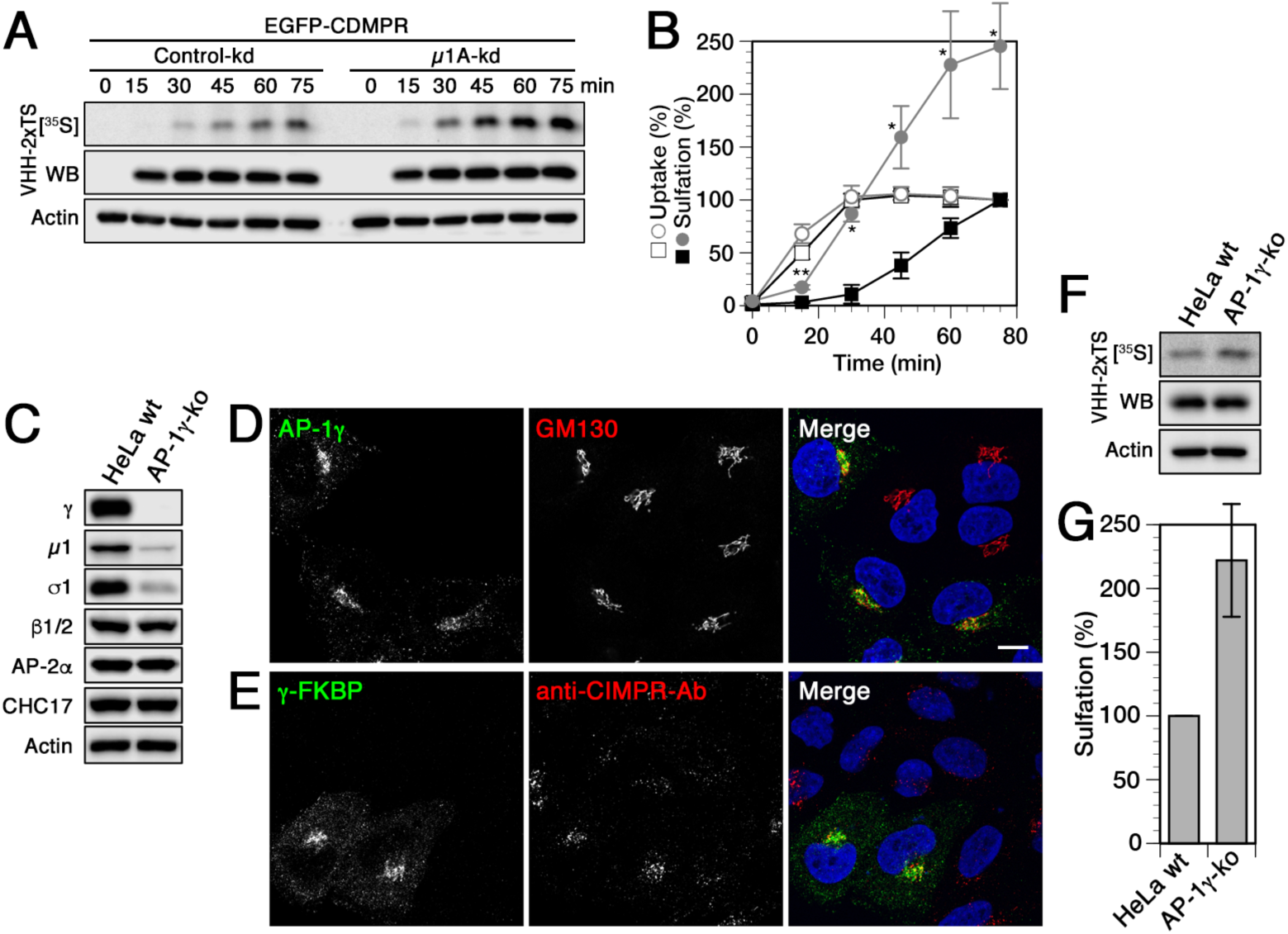
Nanobodies transported to the TGN by CDMPR upon knockdown or knockout of AP-1 is hypersulfated. (**A** and **B**) Cells stably expressing EGFP-CDMPR were transfected with non-targeting siRNA (control-kd) or with siRNA silencing expression of μ1A as described in Fig.1. The cells were labeled with [^35^S]sulfate for up to 75 min in the presence of 2 μg/ml VHH-2xTS and the nanobodies were isolated, analyzed, and quantified as in Fig. 2 (mean and standard deviation of three independent experiments; two-sided Student’s *t*-test: *, p < 0.05; **, p < 0.01). Control-kd is shown as black squares and μ1A-kd as gray circles; uptake as open symbols, sulfation as filled symbols. (**C**) Immunoblot analysis of parental HeLa cells (HeLa wt) and of a pool of γ-adaptin-knockout cells (AP-1γ-ko) generated with CRISPR/Cas9. Equal amounts of cell lysates were probed with antibodies against specific AP-1 subunits (γ, μ1A, σ1), β-adaptins of AP-1 and AP-2 (β1/2), AP-2α, clathrin heavy-chain (CHC17), and actin. (**D**) Parental HeLa cells and AP-1γ knockout cells were mixed and stained with antibodies targeting AP-1γ or GM130. γ-Adaptin staining was completely absent in knockout cells, while Golgi morphology remained intact. (**E**) AP-1γ knockout cells were transiently transfected with γ-FKBP (a fusion protein of γ-adaptin with FK506 binding protein; see Materials and Methods) and incubated with anti-CIMPR antibody for 1 h at 37°C. Cells were fixed and prepared for immunofluorescence microscopy by staining the recombinant γ-subunit and the internalized antibody. Non-transfected cells mostly displayed peripheral accumulation of anti-CIMPR antibody, while expression of γ-FKBP largely rescued perinuclear anti-CIMPR antibody localization. Nuclei were stained with DAPI (blue). Bar: 10 *µ*m. (**F** and **G**) Parental HeLa and AP-1γ knockout cells were transiently transfected with EGFP-CDMPR, followed by [^35^S]sulfate labeling for 75 min in the presence of 2 *µ*g/ml VHH-2xTS. The nanobodies were isolated, analyzed, and quantified as in Fig. 2 (mean and standard deviation of three independent experiments).

To rule out artefacts like off-target effects of siRNA-mediated silencing of μ1A, we generated HeLa AP-1 knockout (ko) cells in which the γ1-adaptin genes were inactivated using CRISPR/Cas9. As expected, knockout cells displayed a complete loss of γ-adaptin staining in immunoblot and immunofluorescence analysis and a concomitant reduction of μ1A and σ1A subunits (Fig. 3C and D). HeLa AP-1γ-ko cells recapitulated the phenotypes of *µ*1A knockdown cells. While Golgi morphology remained unchanged, internalized anti-CIMPR antibody showed mainly peripheral localization that was largely rescued to a predominantly perinuclear Golgi localization upon re-expression of a γ-adaptin fusion protein (Fig. 3E). Just like in the *µ*1A knockdown cells, sulfation of nanobody internalized by transfected EGFP-CDMPR was at least two-fold higher after 75 min of labeling than in wild-type HeLa cells (Fig. 3F and G).

However, we have previously observed that sulfation is not simply a detector of arrival in the compartment. Nanobody sulfation appears not to be sufficiently efficient to immediately and completely modify the sulfation sites as they enter the sulfation compartment. This explains why sulfation per nanobody depended on the target receptor: nanobodies taken up by EGFP-TGN46 showed considerably higher specific sulfation within 1 h than those captured by the EGFP-MPRs, even though maximal nanobody uptake was reached much later (Buser et al., 2018). This most likely reflects the residence time of these proteins in the sulfation compartment during which sulfate was continually incorporated into the nanobodies. A potential explanation of the observed hypersulfation is thus an increased residence time of the nanobody–EGFP-CDMPR complexes that still reached the TGN in the absence of AP-1. This is not unlikely, since AP-1/clathrin is not only involved in retrograde retrieval to the TGN, but also in anterograde transport of MPRs out of the TGN to endosomes (Doray et al., 2002; Ghosh et al., 2003a; Traub et al., 1995; Traub et al., 1993; Waguri et al., 2003).

### Rapid AP-1 inactivation by knocksideways inhibits TGN exit of CDMPR

The bidirectional function of AP-1/clathrin in MPR traffic thus makes it impossible to directly compare nanobody sulfation kinetics with other knockdown situations. To more directly demonstrate the effect of an anterograde transport block at the TGN on nanobody sulfation, we employed the AP-1 knocksideways cells (HeLa-AP1ks) established previously (Buser et al., 2018). AP-1 rerouting to mitochondria by rapamycin for 1 h shifted the steady-state distribution of EGFP-CDMPR to peripheral compartments as expected (Fig. 4A and B). To demonstrate AP-1 dependence of TGN exit, we first loaded HeLa-AP1ks cells expressing EGFP-CDMPR with VHH-2xTS nanobody to steady-state during sulfate starvation, followed by [^35^S]sulfate labeling for up to 75 min. Upon addition of [^35^S]sulfate, there is a delay of 2–3 min for uptake and formation of 3’-phosphoadenosine-5’-phosphosulfate (PAPS) before incorporation of radiactivity gradually starts. To avoid these starting effects, rapamycin was added only 15 min after addition of [^35^S]sulfate. Inactivation of AP-1 caused a more than two-fold increase in sulfation rate compared to cells treated with vehicle only, to reach saturation within the next 15 min, much earlier than in control cells (Fig. 4C-D). While entry of CDMPR into the TGN is reduced by the rapid depletion of available AP-1 as we previously observed in retrograde transport experiment with the same cell line (Buser et al., 2018), the observed increase in sulfation in the present experiment thus reflects the accumulation of nanobody–EGFP-CDMPR in the compartment of sulfation due to reduced TGN exit. This offers itself also as an explanation of the hypersulfation in the TGN-arrival assay of Figure 3.

**Figure 4.**
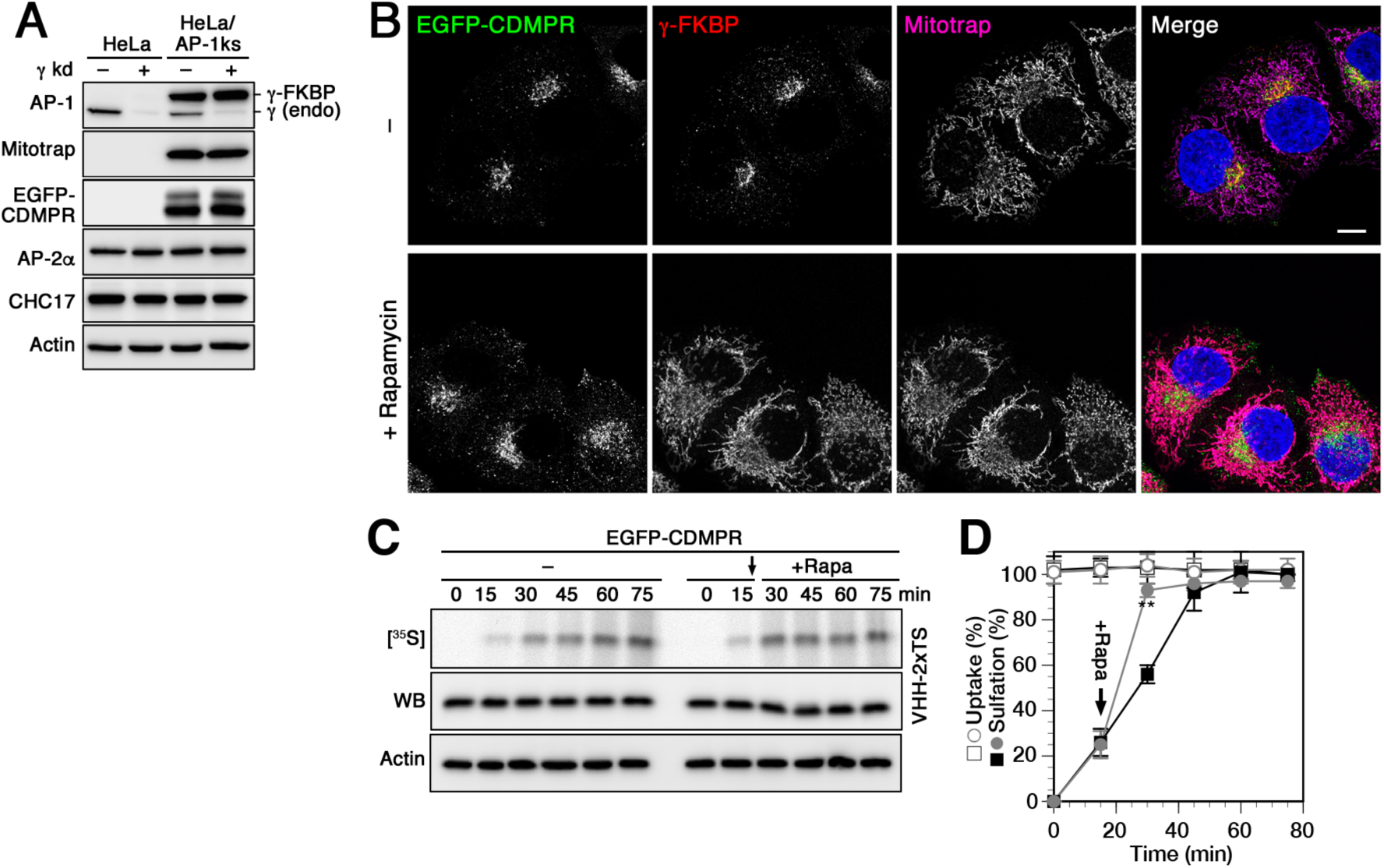
Increased nanobody sulfation upon AP-1 silencing is the consequence of anterograde TGN exit block. (**A**) Lysates of normal HeLa cells and HeLa-AP1 knocksideways (HeLa-AP1ks) cells stably expressing γ-FKBP, Mitotrap and EGFP-CDMPR with or without siRNA-mediated knockdown of the endogenous γ-adaptin were subjected to immunoblot analysis for both forms of γ-adaptin, for Mitotrap (anti-FLAG), EGFP-CDMPR, the α-adaptin subunit of AP-2, and clathrin heavy-chain (CHC17). Knockdown efficiencies for endogenous γ-adaptin were typically >85%. (**B**) HeLa-AP1ks cells stably expressing EGFP-CDMPR after silencing endogenous γ-adaptin were treated with or without 500 nM rapamycin for 1 h and processed for fluorescence microscopy to detect EGFP-CDMPR, recombinant γ-FKBP (using an antibody targeting an epitope present in the neuronal splice variant of AP-2α), and Mitotrap (anti-FLAG). Bar: 10 *µ*m. (**C**) HeLa-AP1ks cells stably expressing EGFP-CDMPR were siRNA-silenced for endogenous γ-adaptin, followed by starvation for sulfate in the presence of VHH-2xTS. The cells were then labeled with [^35^S]sulfate for up to 75 min, without or with addition of 500 nM rapamycin after 15 min (arrow) to inactivate AP-1 (+rapa). The nanobodies were isolated by Ni/NTA beads and subjected to SDS-gel electrophoresis followed by immunoblot analysis (anti-His6) and autoradiography ([^35^S]). In parallel, aliquots of the cell lysates were immunoblotted for actin as a control for the amount of cells used. (**D**) Three independent experiments as shown in panels C were quantified and presented as the percentage of the value in the absence of rapamycin after 75 min (mean and standard deviation of three independent experiments; two-sided Student’s *t*-test: *, p < 0.05; **, p < 0.01). Without rapamycin is shown as black squares, with rapamycin as gray circles; uptake as open symbols, sulfation as filled symbols.

### EpsinR and GGAs depletion both affect retrograde transport to the TGN

The exceptional role of AP-1 in mediating both anterograde and retrograde transport prompted us to consider how other adaptor proteins that at least partially co-operate with AP-1 influence retrograde traffic of CDMPR, in particular epsinR and the GGA adaptors (GGA1–3). In epsinR-depleted cells, isolated CCVs displayed a ∼50% loss of incorporated AP-1, suggesting that AP-1 is to some extent dependent on epsinR for its incorporation into CCVs (Hirst et al., 2004; Hirst et al., 2003). GGAs have been described to play a role in the packaging of MPRs into anterograde AP-1/clathrin carriers at the TGN (Doray et al., 2002; Ghosh and Kornfeld, 2004). This was further supported by the preferential depletion of lysosomal hydrolases and their receptors from CCVs upon GGA2 knocksideways (Hirst et al., 2012).

Levels of GGAs, individually or in combination, and of epsinR could be efficiently reduced by >85% by RNAi, while other sorting machineries remained unperturbed (Fig. 5A and B). Also the steady-state levels of EGFP-CDMPR remained unaffected (Fig. S1). Depleting epsinR in cells stably expressing EGFP-CDMPR did not considerably affect reporter-nanobody localization, with only a slight increase of MPRs redistributed to the periphery (Fig. 5C and D). This observation is in agreement with previous findings showing no effect on the steady-state localization of CIMPR and furin upon silencing of epsinR (Hirst et al., 2004). Reducing GGA levels impacted the localization of MPR–nanobody similarly to the depletion of AP-1 (Fig. 5C and D, compare to Fig. 1E and F), in agreement with previous findings (Ghosh et al., 2003b). This phenotype is intriguing, since it is rather characteristic for proteins mediating retrograde transport.

**Figure 5.**
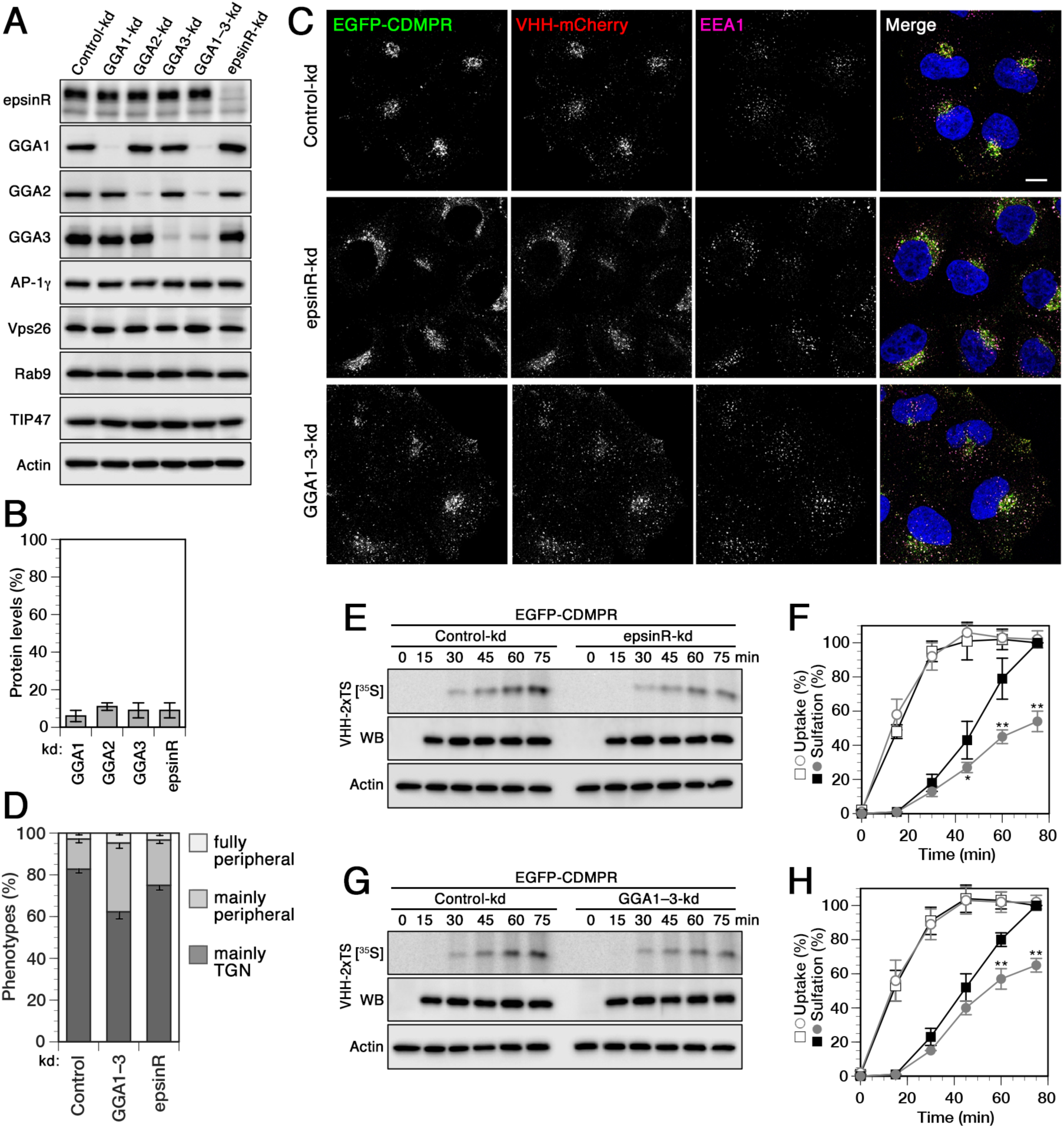
Knockdown of the clathrin adaptors epsinR and GGA1–3 reduces retrograde transport of CDMPR. (**A**) HeLa cells were transfected with non-targeting siRNA (Control-kd) or siRNAs targeting GGA1, GGA2, and GGA3, individually or combined (GGA1–3-kd), or epsinR. Three days after transfection, cells were analyzed by immunoblotting with antibodies against the indicated proteins. (**B**) To determine the knockdown efficiency, the residual protein was quantified in percent of the value after control-kd (mean and standard deviation of three independent experiments). (**C**) HeLa cells stably expressing EGFP-CDMPR were depleted of epsinR, or all three GGAs (GGA1–3) as in (B). Cells were incubated for 1 h at 37°C with full medium containing 5 *µ*g/ml VHH-mCherry (∼0.1 *µ*M), fixed, stained for EEA1 and nuclei (DAPI, blue), and imaged by fluorescence microscopy. Bar: 10 *µ*m. (**D**) Quantitation of the percentage of cells displaying the CDMPR localization phenotypes “mainly TGN”, “mainly peripheral”, or “fully peripheral” (Simonetti et al., 2017; Wassmer et al., 2007). For each condition, random frames with a total of 137–152 cells were scored from three independent experiments. (**E–H**) Cells stably expressing EGFP-CDMPR were transfected with non-targeting siRNA (control-kd) or with siRNA silencing expression of epsinR (E and F) or GGA1–3 (G and H) as described in (A). The cells were labeled with [^35^S]sulfate for up to 75 min in the presence of 2 *µ*g/ml VHH-2xTS and the nanobodies were isolated, analyzed, and quantified as in Fig. 2 (mean and standard deviation of three independent experiments; two-sided Student’s *t*-test: *, p < 0.05; **, p < 0.01). Control-kd is shown as black squares and μ1A-kd as gray circles; uptake as open symbols, sulfation as filled symbols.

Using our nanobody sulfation assay to determine the contributions of epsinR and GGAs on CDMPR transport, one would expect to find a reduction in sulfation kinetics to indicate inhibition of retrograde transport or hypersulfation to indicate inhibition of anterograde TGN exit. EpsinR-depleted cells showed no difference in nanobody uptake, but a strong impairment in retrograde transport of EGFP-CDMPR by ∼50% after 75 min (Fig. 5E-F), comparable to the effect of Vps26 depletion (Fig. 2A-B). This result confirms a role of epsinR in endosome-to-TGN retrograde transport of CDMPR as previously suggested for CIMPR, TGN46, and STxB by Johannes and colleagues (Saint-Pol et al., 2004).

Surprisingly, however, depletion of the GGAs revealed a very similar and significant reduction of VHH-2xTS sulfation kinetics by ∼40% with no apparent effect on uptake (Fig. 5G and H). This result is not consistent with the expected unique function of GGAs in anterograde transport, but rather supports a role in retrograde traffic.

## Discussion

In the present study, we have systematically analyzed the contribution of various intracellular sorting machineries to retrieval of CDMPR from the cell surface to the TGN by a transport assay. Most earlier studies were based on the analysis of changes in steady-state distributions of transported proteins by fluorescence microscopy upon silencing of potential machinery components. To more directly and quantitatively assess transport and its kinetics, we took advantage of functionalized anti-GFP nanobodies in combination with cell lines stably expressing EGFP-CDMPR (Buser et al., 2018; Buser and Spiess, 2019). Nanobodies containing TS sites report arrival and residence in the TGN, the compartment of sulfation, as they are internalized piggyback by EGFP-CDMPR from the cell surface. We previously applied this assay to test the contribution of AP-1 using the knocksideways system for rapid depletion (Buser et al., 2018). Since AP-1 is also implicated in anterograde transport from the TGN to endosomes, rapid inactivation promised less indirect effects resulting from inhibition of TGN exit than long-term silencing by siRNA-mediated knockdown or knockout. A clear reduction of the rate of sulfation by approximately one third was detected after rapamycin-induced knocksideways, thus confirming a non-exclusive role of AP-1/clathrin in retrograde transport of CDMPR.

Here, we performed the experiment also with and without AP-1 inactivation by knockdown or knockout. The result was indeed strikingly different, since hypersulfation was observed (Fig. 3). While other, AP-1-independent pathways still mediate significant retrograde transport of nanobody–EGFP-CDMPR complexes from endosomes to the TGN, their exit from the TGN is reduced by the absence of AP-1, extending their residence time in the sulfation compartment and thus the incorporation of [^35^S]sulfate. We could show that pre-equilibrated nanobody–EGFP-CDMPR at the TGN was more strongly sulfated as soon as AP-1 depletion was triggered by rapamycin addition in knocksideways cells (Fig. 4), clear evidence for inhibition of TGN exit and the anterograde role of AP-1-CCVs for CDMPR. Upon long-term depletion of AP-1, the steady-state pool of CDMPR at the TGN is likely higher than immediately after rapamycin-triggered knocksideways. This will further increase the residence time in the TGN and thus sulfation of entering nanobody–EGFP-CDMPR complexes and may account for the strong increase in sulfation after knockdown and knocksideways that overcompensates the reduction in incoming CDMPR.

It cannot be excluded that additional indirect effects caused by gradual and long-term depletion contribute to hypersulfation. Several unexpected and unexplained phenomena have previously been observed upon knockdown of AP-1, but not upon knocksideways: almost no reduction of CIMPR and ARF1 in CCVs, but increased AP-2 levels (Navarro Negredo et al., 2017; Robinson et al., 2010), and GGA2 was still incorporated into CCVs isolated from AP-1 knockdown but not from knocksideways cells (Hirst et al., 2012). However, tyrosine sulfation activity was not affected in AP-1 knockdown cells (Fig. S2). Interestingly, in a proteomics search for CCV content dependent on AP-1, SLC35B2, one of the transporters delivering the activated sulfation precursor 3’-phosphoadenosine-5’-phosphosulfate (PAPS) into the TGN lumen scored positive (Hirst et al., 2012). One might thus speculate that lack of retrieval of this and other components of the sulfation machinery specifically by AP-1 might lead to hypersulfation in endosomes. While this scenario appears unlikely, one should mention that Johannes and colleagues in fact observed a slight increase in STxB sulfation, when AP-1 was silenced by siRNA (Saint-Pol et al., 2004).

In any case, using our assay, it is expected that silencing of components involved in retrograde transport machineries for CDMPR causes reduced rates of nanobody sulfation, and depletion of proteins mediating TGN exit causes increased rates of sulfation. Accordingly, we found a clear reduction of nanobody sulfation upon knockdown of Vps26, confirming the role of retromer in retrograde transport also for CDMPR. There are many reports for a requirement of retromer for CIMPR (Arighi et al., 2004; Bugarcic et al., 2011; Bulankina et al., 2009; Chen et al., 2019; Cui et al., 2019; Fjorback et al., 2012; Hao et al., 2013; Harbour et al., 2010; Hirst et al., 2018; McGough et al., 2014; Seaman, 2004; Seaman, 2007; Seaman et al., 2009; Seaman, 2018; Wang et al., 2018; Wassmer et al., 2007). However, the requirement of the trimeric Vps retromer complex for CIMPR retrieval has recently been challenged by reports of the Cullen and Steinberg labs (Kvainickas et al., 2017; Simonetti et al., 2017). Using colocalization analyses, they failed to detect mislocalized CIMPR in Vps35-silenced cells, but identified a specific motif in CIMPR’s cytosolic tail (WLM, not present in CDMPR) that binds to the SNX-BARs SNX1/2 and SNX5/6 and is required for correct receptor localization. CI- and CDMPRs may well differ in their interactions with retromer components. Furthermore, loss of either Vps26 or Vps35, two components of the retromer core complex, produced very different phenotypes for retrograde STxB transport to the TGN (Popoff et al., 2009), showing potential alternate functions of the retromer subunits.

Defects in retrograde transport of CDMPR characteristically goes together with a change in steady-state distribution of the receptor in favor of peripheral endosomal compartments. This was found to be the case, when Vps26 or AP-1 was silenced, but not detectably for TIP47 and only to a small extent for Rab9 (Fig. 1). In agreement with this result, no change in nanobody sulfation was observed upon TIP47 knockdown and only a small reduction upon Rab9 silencing (Fig. 2). Our results thus do not confirm a role of TIP47 as a sorting device for CDMPR transport in vivo, but does show an impact of Rab9 depletion on CDMPR arrival in the TGN. The effect of Rab9 depletion on retrograde MPR transport might also be the result of effects on endosomal maturation in general.

Silencing epsinR was found to cause a clear reduction of nanobody sulfation and a slight redistribution of CDMPR to the periphery (Fig. 5), consistent with a role of epsinR in retrograde transport of CDMPR. This result adds to previous studies showing a function of epsinR on distribution or transport of CIMPR and STxB (Saint-Pol et al., 2004).

Most surprising was our finding that depletion of GGA1–3 did not produce hypersulfation of nanobodies imported by EGFP-CDMPR as expected for a component involved in TGN exit, but rather reduced sulfation indicative of a defect in retrograde transport (Fig. 5). Consistent with this notion, significant peripheral redistribution of CDMPR was observed and not perinuclear accumulation at the TGN. The results contradict an exclusive role of GGAs in anterograde transport of MPRs in cooperation with AP-1 as previously proposed (Doray et al., 2002; Ghosh et al., 2003a; Ghosh and Kornfeld, 2004). GGAs clearly localize both to the trans-Golgi and to endosomes (D’Souza et al., 2014; Dell’Angelica et al., 2000; Ghosh et al., 2003b; Hirst et al., 2000; Ratcliffe et al., 2016; Uemura et al., 2018; Wahle et al., 2005), like AP-1, and thus could operate at both places. Since GGA depletion had previously been observed to cause redistribution of CIMPR to EEA1-positive compartments, a role also in retrograde transport from endosomes had not been excluded (Ghosh et al., 2003b). Comparative CCV proteomics with GGA2 and AP-1 knocksideways cells pointed towards involvement of GGA/AP-1 coats in anterograde sorting of MPR–lysosomal hydrolase complexes from the TGN (Hirst et al., 2012), yet the authors did not discount a potential retrograde function. Interestingly, a recent study perfomed in *Schizosacchoromyces pombe* demonstrated that GGAs in collaboration with clathrin adaptors indeed contribute to efficient retrograde transport of Vps10, yeast’s MPR homologue, from the prevaculaor endosome to the TGN (Yanguas et al., 2019). In addition, despite the functional relationship of GGA2 and AP-1 adaptors, it is suprising to observe that they are spatially segregated from each other to a considerable extent (Huang et al., 2019). Our results showing reduced CDMPR transport to the TGN using a sulfation-based approach strongly support a contribution to retrograde transport by GGAs.

In the present study, we have analyzed the contribution of individual sorting machineries in retrograde endosome-to-TGN transport of CDMPR and found that several machineries contribute, likely from different types of endosomes: retromer, the clathrin adaptors AP-1, epsinR, and – most surprisingly – the GGAs, and to some extent Rab9. Other sorting machineries that might facilitate endosome-to-TGN transport of CDMPR include SNX-BAR proteins and AP-5 (Hirst et al., 2018; Kvainickas et al., 2017; Simonetti et al., 2017; Simonetti et al., 2019). Both have been shown to be important for CIMPR retrieval, not yet for CDMPR. Future systematic and comparative analysis of machinery requirement between CDMPR and CIMPR in retrograde transport for these machineries are important to further understand the coexistence and operation of multiple TGN retrieval pathways for receptors. Our approach of sulfatable nanobodies offers new avenues in understanding sorting machinery contribution for cargo proteins of interest.

## Materials and Methods

### Bacterial expression and purification of functionalized nanobodies

Functionalized nanobodies were bacterially expressed and isolated as previously described (Buser et al., 2018; Buser and Spiess, 2019). Briefly, bacterial expression vectors encoding derivatized VHH nanobodies and myc-BirA (Addgene #109424) were transformed together into Rosetta DE3 cells (Merck), and plated on LB plates with 50 *µ*g/ml kanamycin and 50 *µ*g/ml carbenicillin. A 20-ml overnight culture of a single colony was diluted into 1 l LB medium with antibiotics and 200 *µ*M D-biotin and grown to an OD_600_ of 0.6-0.7 at 37°C. Expression was induced with 1 mM isopropyl-β-D-thiogalactopyranosid (IPTG) at 16°C overnight (VHH-mCherry), or at 30°C for 4 h (VHH-std and VHH-2xTS). Cells were pelleted at 5’000×g at 4°C for 45 min and stored at –80°C. Upon thawing, they were resuspended in 30 ml PBS with 20 mM imidazole, 200 *µ*g/ml lysozyme, 20 *µ*g/ml DNase I, 1 mM MgCl_2_, and 1 mM PMSF, incubated for 10 min at room temperature and 1 h at 4°C while rotating, followed by mechanical lysis using a tip sonicator for 3 times 30 s with 1-min cooling periods. The lysate was cleared by centrifugation at 15’000×g for 1 h at 4°C and loaded on a His GraviTrap column (GE Healthcare Life Sciences), washed with 20 mM imidazole in PBS, and eluted with 2 ml PBS with 500 mM imidazole. The purified nanobodies were desalted on PD-10 columns (GE Healthcare Life Sciences), concentrated to 2 mg/ml (for VHH-std and VHH-2xTS) or 5 mg/ml (for VHH-mCherry), flash-frozen in liquid nitrogen, and stored at –80°C. Plasmids for nanobody fusion protein expression are deposited with Addgene (Addgene: #109417, VHH-std; #109419, VHH-2xTS; #109421, VHH-mCherry).

### Cell culture, CRISPR/Cas9 gene editing, and RNA interference

HeLa α cell lines were maintained in high-glucose Dulbecco’s modified Eagle’s medium (DMEM) with 10% fetal calf serum (FCS), 100 units/ml streptomycin, 2 mM L-glutamine and appropriate selection antibiotics (1.5 *µ*g/ml puromycin, 1 mg/ml hygromycin B, or 7.5 *µ*g/ml blasticidin) at 37°C in 7.5% CO_2_. Phoenix Ampho packaging cells (from the Nolan lab, Stanford University) were grown in complete medium supplemented with 1 mM sodium pyruvate.

HeLa cells stably expressing EGFP-CDMPR, and HeLa-AP1ks cells stably expressing the respective EGFP reporter were previously described (Buser et al., 2018). To generate HeLa cells stably expressing sulfatable A1Pi, Phoenix Ampho packaging cells were transfected pQCXIP-SHMY-A1Pi using FuGENE HD (Promega). The viral supernatant was harvested after 48 h, passed through a 0.45 *µ*m filter, supplemented with 15 *µ*g/ml polybrene, and added to target HeLa α cells. The next day, complete medium with 1.5 *µ*g/ml puromycin was added for selection, and pooled resistant clones were used for experiments.

To generate a γ-adaptin knockout HeLa cell line by CRISPR/Cas9, sgRNAs for gene editing were purchased from Santa Cruz Biotechnology (sc-403986). Briefly, parental HeLa cells were transfected with 2 μg plasmid containing a GFP cassette and 4 μl FuGENE HD (Promega) in a six-well cluster. After 24 h of expression, cells were subjected to fluorescence-activated cell sorting (FACS), and single cells or pooled clones were collected. Transient transfection of AP-1ko cells with pQCXIP-γ-FKBP was performed using FuGENE HD according to the manufacturer’s instructions. Anti-CIMPR antibody uptake was performed as described previously (Robinson et al., 2010).

For RNA interference, cells were reverse-transfected with target siRNA in Opti-MEM I using Lipofectamine RNAiMAX (both Thermo Fisher Scientific) following the manufacturer’s instructions. For a conventional knockdown of AP-1, the sequence 5’-AAGGCAUCAAGUAUCGGAAGAdTdT-3’ against the μ1A-subunit of the heterotetrameric complex was used as formerly reported (Hirst et al., 2005; Hirst et al., 2003; Hirst et al., 2009). For RNA interference with retromer complex, we applied siRNA duplexes with the sequence 5’- AACUCCUGUAACCCUUGAGdTdT-3’ targeting Vps26 as described in previous studies (Popoff et al., 2009; Popoff et al., 2007). To specifically silence Rab9 or TIP47, we applied the siRNA sequence 5’- GUUUGAUACCCAGCUCUUCdTdT 3’ for Rab9 (Ganley et al., 2004; Kucera et al., 2016; Reddy et al., 2006), or 5’-CCCGGGGCUCAUUUCAAACdTdT-3’ for TIP47 (Bulankina et al., 2009). EpsinR was targeted with 5’- AAUACAGAUAUGGUCCAGAAATTdTdT-3’, GGA1 with 5’-CACAGGAGUGGGAGGCGAUTTdTdT-3’, GGA2 with 5’-UGAAUUAUGUUUCGCAGAATTdTdT-3’, and GGA3 with 5’- UGUGACAGCCUACGAUAAATTdTdT-3’ as previously described (Hirst et al., 2012; Hirst et al., 2004). To knockdown AP-2α and CHC17, we used 5’-AAGAGCAUGUGCACGCUGGCCAdTdT-3’ and 5’- UAAUCCAAUUCGAAGACCAAUdTdT-3’ duplexes, respectively, as previously described (Motley et al., 2003). We used the non-targeting siRNA 5’-UAAGGCUAUGAAGAGAUACdTdT-3’ as control siRNA (Salazar et al., 2009). All siRNAs were used at a final concentration of 50 nM, apart from the GGA siRNAs (used at 25 nM each).

To silence γ-adaptin in HeLa-AP1ks cells, the siRNA sequence 5’-GAAGAUAGAAUUCACCUUUUU-3’ was used as previously described (Buser et al., 2018; Robinson et al., 2010). Cells were transfected twice (day 1 and 3) and used at day 5. siRNA duplexes were purchased from Microsynth.

### Gel electrophoresis and immunoblot analysis

Proteins separated by SDS-gel electrophoresis (7.5–15% polyacrylamide) were transferred to Immobilon-P^SQ^ PVDF membranes (Millipore). After blocking with 5% non-fat dry milk in TBS (50 mM Tris·HCl, pH 7.6, 150 mM NaCl) with 0.1% Tween-20 (TBST) for 1 h, the membranes were probed with primary antibodies in 1% BSA in TBST for 2 h at room temperature or overnight at 4°C, followed by incubation with HRP-coupled secondary antibodies in 1% BSA in TBST for 1 h at room temperature. Immobilon Western Chemiluminescent HRP Substrate (Millipore) was used for detection, a Fusion Vilber Lourmat Imaging System for imaging, and Fiji software for quantitation.

### Fluorescence microscopy

For immunofluorescence staining, cells were grown on glass coverslips, fixed with 3% paraformaldehyde (PFA) for 10 min at room temperature, washed with PBS, quenched with 50 mM NH_4_Cl in PBS for 5 min, permeabilized with 0.1% Triton X-100 in PBS for 10 min, blocked with 1% BSA in PBS for at least 15 min, incubated with primary antibody in BSA/PBS for 2 h, washed, and stained with fluorescent secondary antibodies in BSA/PBS for 1 h. After a 5 min staining with 5 μg/ml DAPI and three washes with PBS, coverslips were mounted in Fluoromount-G (Southern Biotech). Staining patterns were imaged on a Zeiss Point Scanning Confocal LSM700 microscope. Receptor mislocalization analysis was performed by scoring and quantification of the percentage of cells displaying each phenotype in machinery-depleted cells using a Zeiss Axioplan microscope with a Leica DFC420C imaging system as described previously (Simonetti et al., 2017; Wassmer et al., 2007).

### Sulfation analysis

To analyze retrograde transport and kinetics of EGFP-CDMPR to the compartment of sulfation, cells were incubated with 1 ml sulfate-free medium for 1 h at 37°C and 7.5% CO_2_ before labeling with sulfate-free medium supplemented with 0.5 mCi/ml [^35^S]sulfate (Hartmann Analytics) and 2 μg/ml purified VHH-2xTS for up to 75 min. For the knocksideways experiment, HeLa-AP1ks cells stably expressing EGFP-CDMPR were starved with sulfate-free medium for 1 h at 37°C and 7.5% CO_2_ in the presence of 2 μg/ml purified VHH-2xTS, followed by two 1 ml washes with sulfate-free medium, prior to labeling with medium reconstituted with 0.5 mCi/ml [^35^S]sulfate. 500 nM rapamycin from a 2000x stock solution in DMSO, or DMSO alone was added. Cells stably expressing SHMY-A1Pi were starved as described above and pulsed with medium reconstituted with 0.5 mCi/ml [^35^S]sulfate for 75 min.

After incubation, cells were washed twice with ice-cold PBS, lysed in 1 ml lysis buffer containing 2 mM PMSF and protease inhibitor cocktail, and centrifuged at 10’000×g for 15 min at 4°C. A fraction (50–100 *µ*l) of the postnuclear supernatants was used for immunoblot analysis of total cell-associated nanobody and an actin control. The rest was incubated for 1 h at 4°C with 20 *µ*l Ni Sepharose High Performance beads (GE Healthcare Life Sciences) to isolate the nanobodies. Beads were washed three times with lysis buffer containing 20 mM imidazole and boiled in SDS-sample buffer. Nanobodies were analyzed by SDS-gel electrophoresis and autoradiography using BAS Storage Phosphor Screens and a Typhoon FLA7000 IP phosphorimager (GE Healthcare Life Sciences).

### Antibodies

For immunofluorescence microscopy, goat anti-α-adaptin (Everest Biotech #EB11875; 1:1000), mouse anti-CIMPR (Abcam ab2733; 1:5000), anti-EEA1 (BD Biosciences #610456; 1:1000), rabbit anti-FLAG (Cell Signaling Technology #2368; 1:500), and rabbit anti-GM130 (Cell Signaling Technology #12480; 1:1000) antibodies were used.

For immunoblotting, mouse anti-α-adaptin (BD Biosciences #610501; 1:5000), mouse anti-γ-adaptin (BD Biosciences #610385; 1:5000), mouse anti-β1/2-adaptin (BD Biosciences #610381; 1:5000), mouse anti-γ-adaptin (made from 100/3 hybridoma; 1:5000), rabbit anti-σ1-adaptin (Bethyl Laboratories #A305-396A-M; 1:1000), mouse anti-γ-adaptin (made from 100/3 hybridoma; 1:5000), mouse anti-actin (EMD Millipore #MAB1501; 1:100000), mouse anti-CHC17 (made from TD.1 hybridoma; 1:200), rabbit anti-epsinR (Bethyl Laboratories #A301-926A; 1:1000), mouse anti-FLAG (Cell Signaling Technology #8146; 1:1000), mouse anti-GFP (Sigma-Aldrich #11814460001-Roche; 1:5000), rabbit anti-GGA1 (Bethyl Laboratories #A305-368A; 1:1000), mouse anti-GGA2 (BD Biosciences #612612; 1:2000), mouse anti-GGA3 (BD Biosciences #612310; 1:1000), mouse anti-HA (made from 12CA5 hybridoma; 1:10000), rabbit anti-His6 (Bethyl Laboratories #A190-114A; 1:10000), rabbit anti-mCherry (GeneTex #GTX128508; 1:10000), rabbit anti-T7 (Bethyl Laboratories #A190-117A; 1:10000), rabbit anti-Rab9 (Cell Signaling Technology #5118; 1:1000), mouse anti-SNX1 (BD Biosciences #611482; 1:500), rabbit anti-SNX2 (Bethyl Laboratories #A304-544A; 1:2000), rabbit anti-TIP47 (Proteintech 10694-1-AP; 1:1000), rabbit anti-Vps26 (Bethyl Laboratories #A304-801A; 1:1000), rabbit anti-Vps35 (Bethyl Laboratories #A304-727A; 1:1000), mouse anti-μ1A (Abnova H00008907-A01; 1:1000) antibodies were used.

As secondary antibodies for immunofluorescence microscopy, A568-labeled donkey anti-goat (Thermo Fisher Scientific #A-11057; 1:500), A633-labeled goat anti-mouse (Thermo Fisher Scientific #A-21052; 1:500), and A633-labeled goat anti-rabbit (Thermo Fisher Scientific #A-21071; 1:500) immunoglobulin antibodies were used. As secondary antibodies for immunoblotting, HRP-labeled goat anti-rabbit (Sigma-Aldrich #A-0545; 1:10000), and goat anti-mouse (Sigma-Aldrich #A-0168; 1:10000) immunoglobulin antibodies were used. To detect biotinylated proteins on blots, Streptavidin-HRP (Thermo Fisher Scientific #434323; 1:10000) was used.

## Acknowledgements

This work was supported by grant 31003A-182519 from the Swiss National Science Foundation. We particularly thank Nicole Beuret, Janine Bögli from the Biozentrum FACS Core Facility, Kai D. Schleicher and Alexia Loynton-Ferrand from the Biozentrum Imaging Core Facility, for their support.

## Author contribution

DPB: planned and carried out experiments, analyzed data, and wrote the manuscript. MS: planned experiments, analyzed data, and wrote the manuscript.

## Figures and Legends

**Figure S1.**
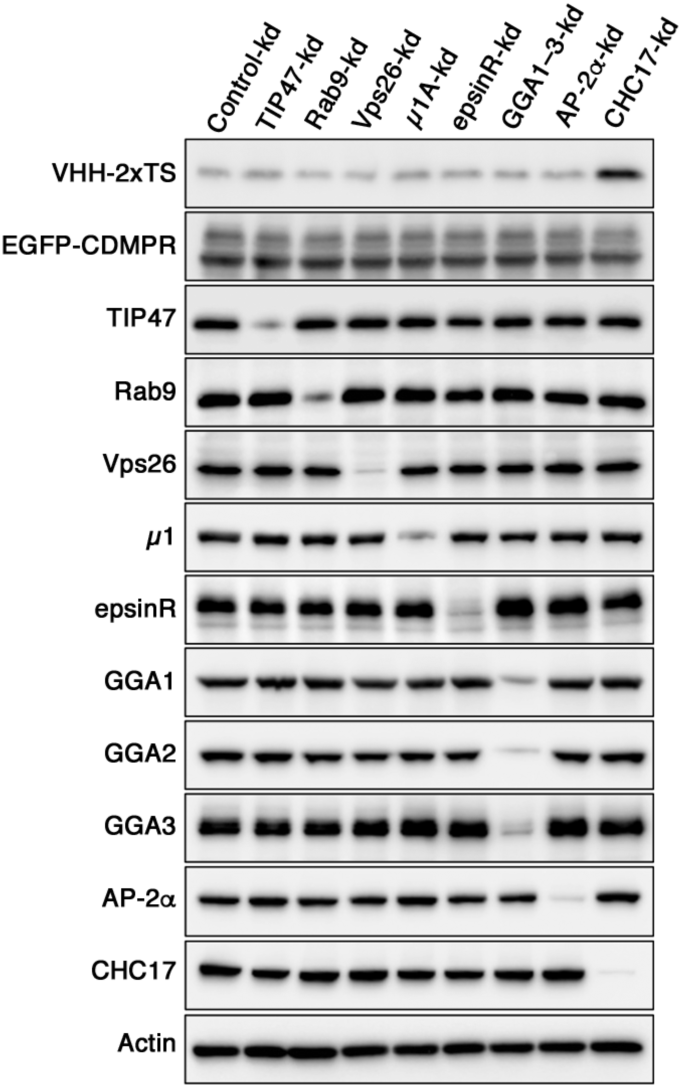
Cell surface levels of EGFP-CDMPR is not altered by silencing of retrograde machinery candidates. HeLa cells stably expressing EGFP-CDMPR were transiently transfected with non-targeting siRNA (Control-kd) or siRNAs targeting TIP47, Rab9, Vps26, μ1A, epsinR, GGA1–3, AP-2α, or clathrin CHC17. Three days after transfection, cells were incubated with 2 μg/ml VHH-2xTS in PBS at 4°C for 1 h to label the surface fraction of EGFP-CDMPR. Subsequently, cells were washed and lyzed, and endogenous protein levels were analyzed by immunoblotting with antibodies against the indicated proteins (VHH-2xTS was detected with anti-His6). Only knockdown of CHC17 influenced surface levels of EGFP-CDMPR (monitored by bound nanobody). Total EGFP-CDMPR remained unaffected in all knockdowns.

**Figure S2.**
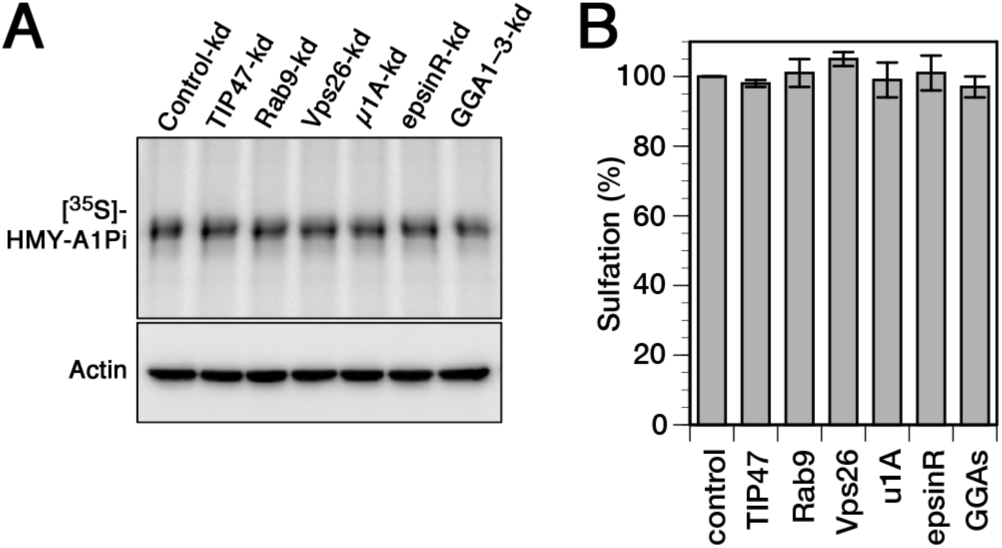
General tyrosine sulfation is not affected by silencing of retrograde machinery candidates. (**A**) HeLa cells stably expressing the sulfatable secretory protein α1-protease inhibitor (or anti-trypsin; SHMY-A1Pi; S, signal sequence of hemagglutinin; H, His6 epitope; M, myc epitope; Y, TS site) were transiently depleted of TIP47, Rab9, Vps26, μ1A-adaptin, epsinR, and GGAs (GGA1–3) for 72 h. Cells were labeled with [^35^S]sulfate labeling for 75 min, lysed, HMY-A1Pi was isolated by Ni/NTA beads, and subjected to SDS-gel electrophoresis followed by autoradiography ([^35^S]). In parallel, aliquots of the cell lysates were immunoblotted for actin as a control for the amount of cells used. (**B**) Quantitation of total tyrosine sulfation as judged by sulfation of the secretory reporter HMY-A1Pi as in panel (A) is shown in percent of the value present in control-kd (mean and standard deviation of three independent experiments).

